# The impact of ischemic stroke on connectivity gradients

**DOI:** 10.1101/481689

**Authors:** Şeyma Bayrak, Ahmed A. Khalil, Kersten Villringer, Jochen B. Fiebach, Arno Villringer, Daniel S. Margulies, Smadar Ovadia-Caro

## Abstract

Understanding the relationship between localized anatomical damage, reorganization, and functional deficits is a major challenge in stroke research. Previous work has shown that localized lesions cause widespread functional connectivity alterations in structurally intact areas, thereby affecting a whole network of interconnected regions. Recent advances suggest an alternative to discrete functional networks by describing a connectivity space based on a low-dimensional embedding of the full connectivity matrix. The dimensions of this space, described as *connectivity gradients*, capture the similarity of areas’ connections along a continuous space. Here, we defined a three-dimensional connectivity space template based on functional connectivity data from healthy controls. By projecting lesion locations into this space, we demonstrate that ischemic strokes resulted in dimension-specific alterations in functional connectivity over the first week after symptoms onset. Specifically, changes in functional connectivity were captured along connectivity Gradients 1 and 3. The degree of change in functional connectivity was determined by the distance from the lesion along these connectivity gradients regardless of the anatomical distance from the lesion. Together, these results provide a novel framework to study reorganization after stroke and suggest that, rather than only impacting on anatomically proximate areas, the indirect effects of ischemic strokes spread along the brain relative to the space defined by its connectivity.

## 1.1 Introduction

Stroke is defined as a sudden neurological deficit caused by a localized injury to the central nervous system due to vascular pathology (Sacco et al., 2013). Outside of the localized structural damage, areas connected to the lesion undergo functional alterations that are implicated in symptomology and the recovery from neurological deficits. This phenomenon is known as *diaschisis* (Andrews, 1991; Carrera and Tononi, 2014) and provides a theoretical and empirical motivation to study brain connectivity following stroke.

Functional connectivity based on the temporal correlation of ongoing blood-oxygen-level-dependent (BOLD) fluctuations (resting-state functional magnetic resonance imaging; rs-fMRI) has been successfully used to study alterations associated with reorganization within functional networks. Previous studies found a reduction in functional connectivity after stroke in structurally intact areas connected to the lesion (i.e., the affected network). Reduction in functional connectivity was associated with the severity of the clinical deficit and recovery of symptoms (Baldassarre et al., 2014; Carter et al., 2010; He et al., 2007; Ovadia-Caro et al., 2013; Siegel et al., 2016; Wang et al., 2010; Warren et al., 2009). Importantly, normalization of connectivity patterns was found following both spontaneous recovery (He et al., 2007; Park et al., 2011; Ramsey et al., 2016; van Meer et al., 2010) and interventions using non-invasive brain stimulation (Volz et al., 2016). Taken together, these findings support the phenomenon of *diaschisis* and the view of stroke as a *network disruption* rather than a mere localized phenomenon (Corbetta, 2010; Ovadia-Caro et al., 2014; Ward, 2005).

While previous studies demonstrate the role of the affected network in stroke pathology, the impact of a lesion is not necessarily limited by network definitions. Graph models of brain connectivity have demonstrated that the local disruption of a single node is likely to extend beyond the affected network and impact, to varying degrees, the whole graph (Aerts et al., 2016; Bassett and Bullmore, 2006; van den Heuvel and Sporns, 2013). Using predefined functional networks assumes sharp boundaries between different functional domains. In addition, it assumes that the effects of stroke are uniformly distributed within a given network. Contrary to these assumptions, recent studies report that connectivity may be better captured by dimensions representing the continuous space of the connectome (Atasoy et al., 2016; Cerliani et al., 2012; Haak et al., 2018). With the shift in our understanding of cognitive brain functions as emerging from global states (Bertolero et al., 2018; Cole et al., 2014; Sporns et al., 2005), so too our models of brain dysfunction should attempt to characterize alterations at the whole-brain level, taking the full connectome into account (see Figure 1).

**Figure 1.**
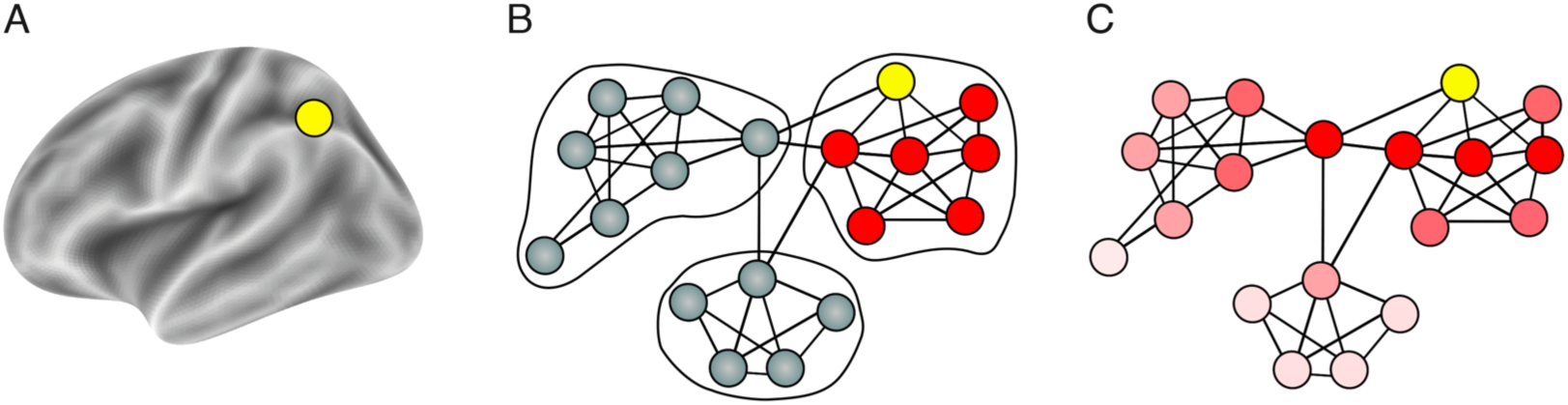
Two complementary views on brain organization and the corresponding representation of distal effects of focal lesions. (A) Representing a focal lesion (yellow node) on the brain anatomical surface. (B) A schematic description of discrete networks parcellation superimposed on a functional connectivity graph-space with nodes and edges. Using this approach to study the effects of focal lesions (yellow node) restricts us to singular networks. Additionally, distal effects of the lesion are assumed to be equally disruptive for all nodes in the affected network (red nodes). (C) Representing functional connectivity in a continuous manner without sharply defined borders using connectivity gradients. The lesioned node affects all other nodes in the system as a function of the distance from the lesion in graph space (dark red to light red). Using this approach does not assume sharp boundaries between functional networks and provides a more realistic model of distant effects of localized lesions.

Recently, non-linear decomposition approaches have been introduced to represent whole-brain rs-fMRI connectivity data in a continuous, low-dimensional space. This data-driven analysis results in *connectivity gradients* that provide a low-dimensional description of the connectome (Langs et al., 2016, 2014; Margulies et al., 2016). Each voxel is located along a connectivity gradient according to its similarity of connections. Voxels that share a similar pattern of functional connectivity are situated close to one another along a given connectivity gradient (Huntenburg et al., 2018). Different functional modules are therefore clustered along a continuum of a given connectivity gradient (Krienen and Sherwood, 2017) without the need of a priori defined network parcellation.

Here, we studied the impact of localized lesions on continuous connectivity gradients. Longitudinal rs-fMRI data were collected from patients following ischemic stroke. Data were collected within 24 hours, as well as one and five days after the onset of stroke symptoms. Changes in functional connectivity over the week were quantified using spatial concordance (Lohmann et al., 2012). Data from healthy subjects were used to create a template of three connectivity gradients representing all possible connections in a continuous manner.

Based on previous findings in discrete networks (Baldassarre et al., 2014; Carter et al., 2010; He et al., 2007; Nomura et al., 2010; Ovadia-Caro et al., 2013; Siegel et al., 2016; Wang et al., 2010; Warren et al., 2009) and computational models (Alstott et al., 2009; Honey and Sporns, 2008; van Dellen et al., 2013; Young et al., 2000), we hypothesized that a lesion along a connectivity gradient would induce a gradual impact on the whole connectome. Functional connectivity alterations would be most pronounced in areas that share a similar connectivity pattern with the lesion.

## 2.1 Materials and methods

### 2.2 Participants

Fifty-four stroke patients (20 females, age: 63.78 ±12.03 years, mean ±SD) and 31 healthy controls (13 females, age: 64.90 ±8.49 years) were initially recruited for the study. Inclusion criteria for patients were: patients older than 18 years, first ever ischemic stroke – small cortical (≤1.5 cm) or subcortical, which was evident in imaging. A Wahlund score ≤ 10 (Wahlund et al., 2001) to limit the extent of white matter lesions. Exclusion criteria included: clinical evidence for antecedent lesions (n=3), fewer than 3 resting-state scans post-stroke (n=10), lesions located solely within white matter (n=3 patients), corrupted MRI raw data or distorted images (n=1 control, n=4 patients), high degree of head motion (n=1 control, n=6 patients), and poor registration quality (n=1 control). For further details on quality assessment see Supplementary Material M1.

Following the exclusion procedure, 28 stroke patients (11 females, age: 65.04 ±13.27 years, mean ±SD), and 28 healthy controls (13 females, age: 65.21 ±8.84 years) were included in the analysis. The groups were matched for age and sex (age: Welch’s t-test, P=0.95; sex: Kruskal-Wallis H-test, P=0.59). For further details on patients’ information see Supplementary Table 1. The study was approved by the ethics committee of the Charité - Universitätsmedizin Berlin, Germany (EA 1/200/13). Written informed consent was obtained from all participants.

### 2.3 Neuroimaging data

The MRI protocol included T1-weighted structural scans and T2*-weighted resting-state fMRI scans (continuous fMRI scan with no overt task) for all participants. In addition, diffusion weighted images (DWI; TR=8.2 s, TE=0.1 s, 50 volumes, voxel size: 2×2×2.5 mm, flip angle 90°) and fluid attenuated inversion recovery images (FLAIR; TR=8.0 s, TE=0.1 s, 54 volumes, voxel size: 0.5×0.5×5 mm) were acquired from the stroke patients as part of a standard MRI protocol (Hotter et al., 2009). All MRI data were acquired on a Siemens Tim Trio 3T scanner. Healthy control participants were scanned at a single time point, whereas stroke patients were scanned at three consecutive time points relative to stroke symptoms onset: day 0 (within 24 hours), day 1 (24 - 48 hours), and day 5 (range: day 4 – 6, mean 4.93 ±0.38 SD). Structural scans were acquired using a three-dimensional magnetization prepared rapid gradient-echo (MPRAGE) sequence (TR=1.9 s, TE=2.52 s, TI=0.9 s, 192 slices, voxel size: 1×1×1 mm, flip angle 9°). Resting-state functional scans for each participant and session were acquired using blood-oxygenation-level-dependent (BOLD) contrast with an EPI sequence (TR=2.3 s, TE=0.03 s, 34 slices, 150 volumes, voxel size: 3×3×3 mm, flip angle 90°, total duration=5.75 min).

### 2.4 Data preprocessing

T1-weighted structural images were preprocessed using FreeSurfer’s recon-all pipeline (v6.0.0, (Dale et al., 1999)). The pipeline generated segmentations for grey matter, white matter and cerebrospinal fluid. Individual grey matter masks were registered to standard MNI space (3 mm^3^).

Preprocessing of functional images included: *i)* removal of the first 5 EPI volumes to avoid signal saturation, *ii)* slice timing and motion correction (Nipype v0.14.0, (Gorgolewski et al., 2011; Roche, 2011)), *iii)* CompCor denoising approach for time series at the voxel level (Nilearn v0.4.0, (Behzadi et al., 2007)), *iv)* temporal normalization, *v)* band-pass filtering in the range of 0.01 - 0.1 Hz, and *vi)* spatial smoothing (applied after registration) with a 6 mm full-width-half maximum Gaussian kernel using FSL (v5.0.9, (Woolrich et al., 2009)). Confounds removed from the time series at the denoising step were defined as *i)* six head motion parameters, including 1st and 2nd order derivatives, *ii)* motion and intensity outliers (Nipype’s rapidart algorithm; thresholds: > 1mm framewise head displacement, and signal intensity > 3 SD of global brain signal accordingly) and *iii)* signal from white matter and cerebrospinal fluid.

The transformation of functional images to MNI152 (3 mm^3^) space included a linear transformation from EPI to the high-resolution T1-weighted image using FreeSurfer’s boundary-based register tool with 6 degrees of freedom (Greve and Fischl, 2009) and a nonlinear transformation using ANTs (v2.1.0, (Avants et al., 2011)). The transformation matrices obtained from both steps were concatenated and applied to the functional image using a single interpolation.

### 2.5 Lesion delineation

Lesions were manually delineated by identifying areas of localized hyperintensity on day 0 DWI images using the ITK-SNAP software (v3.4.0, (Yushkevich et al., 2006)). Delineations were guided by expert radiology reports and were approved by a radiology resident. All lesion masks were normalized to MNI152 (3 mm^3^) space (ANTs, nearest-neighbor interpolation). Individual lesion masks were smoothed in the atlas space using FSL’s dilation tool with 3×3×3 kernel, extending the mask by one voxel-size (v5.0.9, (Jenkinson et al., 2012)).

### 2.6 Computing connectivity gradients by applying nonlinear decomposition to functional connectivity data from healthy controls

To create a mutual grey matter template to be used for decomposition analysis, individual grey matter masks and resting-state functional masks were averaged for all healthy controls to create a group mask. Averaged group maps were multiplied to create a mutual mask such that only grey matter voxels with fMRI signal would be included. The resulting template (33,327 voxels) was used to generate functional connectivity matrices from individual healthy controls.

Functional connectivity matrices (33,327×33,327 voxels) were computed using Pearson’s correlation coefficient and were normalized using Fisher’s z-transformation. An average functional connectivity matrix was computed across healthy controls and the averaged z-scores were transformed back to r-scores. Each row of the group-level functional connectivity matrix was thresholded at 90% of its r-scores. This yielded an asymmetric, sparse matrix. The pairwise cosine similarities of all rows were computed. By doing this, we obtained a non-negative and symmetric similarity matrix, *L* (values in [0, 1] range).

We implemented the diffusion embedding approach on the similarity matrix to obtain a low-dimensional representation of the whole-brain functional connectivity matrix (Coifman and Lafon, 2006; Langs et al., 2016), as done in Margulies et al., 2016. This approach resulted in gradients of functional connectivity. Voxels along each gradient are assigned unitless embedding values. Along each gradient, voxels that share similar connectivity pattern have similar embedding values.

### 2.7 Mapping individual stroke lesions onto connectivity gradients from healthy controls

Individual lesion masks were projected onto the individual gradients obtained in healthy controls. Lesioned voxels were marked according to their location along a specific gradient. The lesion site along each gradient was defined as the minimum embedding value of all lesioned voxels.

To quantify the functional similarity of non-lesioned voxels to the lesion site, distance-to-lesion maps were computed for each non-lesioned voxel (Figure 2B). Distance values reflect the mutual difference between embedding values of non-lesioned and lesioned voxels. Low distance values reflect voxels that share similar functional connectivity pattern with the lesion site.

**Figure 2.**
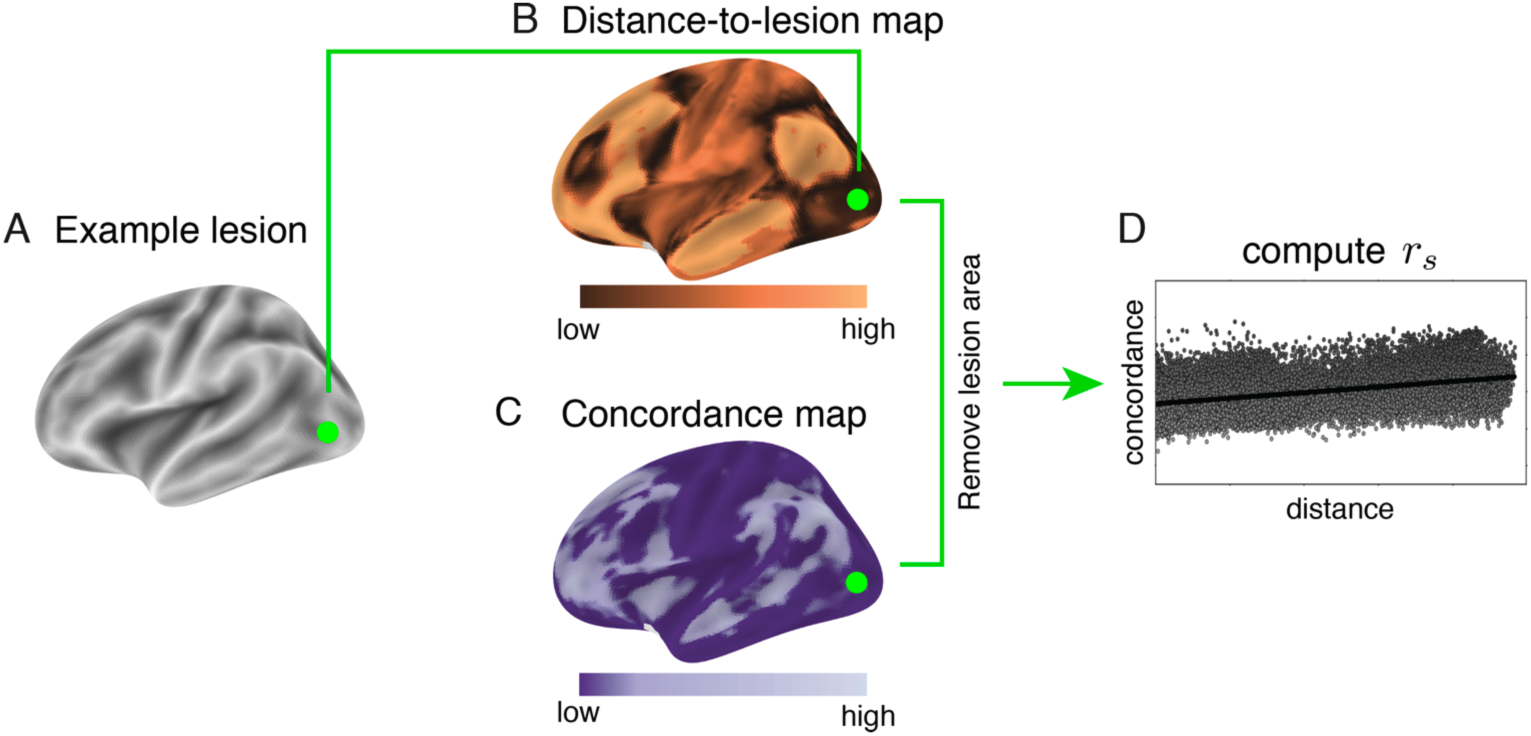
A schematic description of the analysis steps. (A) Individual lesions were delineated for each patient. Here, an example of a lesion located in the left occipital lobe (green). (B) Distance-to-lesion maps were computed for each of the three connectivity gradients. Distance values reflect the mutual difference between embedding values of non-lesioned and lesioned voxels. Low distances (dark-copper) represent voxels that share a similar functional connectivity pattern with the lesion site. This example shows the distance-to-lesion map for the first gradient. (C) A voxel-wise spatial concordance map was computed for each patient across the three resting-state scans after stroke. Concordance correlation coefficient (CCC) values reflect the degree of change in the connectivity pattern over time for each voxel. Low CCC values (dark-purple) represent voxels that underwent a larger change in their functional connectivity pattern over time. (D) Spearman’s rank correlation coefficient (*r*_s_) was used to test the relationship between distance-to-lesion and degree of functional connectivity alteration across all voxels. A positive correlation depicts a larger change in functional connectivity for voxels that were closer to the lesion site along the corresponding connectivity gradient.

### 2.8 Quantifying longitudinal alterations in functional connectivity matrices for stroke patients

For each patient, a functional mask was obtained from each of the three consecutive functional scans. These masks were multiplied with the grey matter template of the healthy cohort. The dilated lesion segmentations were then excluded from the patient-specific grey matter template. This approach ensured that functional images of patients included only identical grey matter voxels as healthy controls, except for the lesion site. The patient-specific grey matter templates varied slightly in number of voxels included (ranging from 32,659 to 33,212 voxels).

To control for the slight variation in the number of voxels in patient-specific grey matter templates, a control analysis was applied such that the grey matter template used for the analysis contained 30,314 voxels in all patients prior to lesion removal. Using this more restricted mask had no influence on our main results (see Supplementary Material M2 and Supplementary Figure S1).

Functional connectivity matrices were computed using Pearson’s correlation coefficient at each of the three time points for individual patients. The voxel-wise spatial concordance map was computed using the concordance correlation coefficient (CCC) (Lin, 2016) at the single-voxel level across the three time points (Lohmann et al., 2012). CCC-values range between -1 and 1, such that the lower concordance reflects larger alterations in the functional connectivity pattern over time (Figure 2C).

### 2.9 The relationship between lesion location along connectivity gradients and alterations in functional connectivity after stroke

Concordance correlation coefficient (CCC) values were correlated with distance-to-lesion values using Spearman’s rank-order correlation coefficient (Figure 2D). This analysis was repeated for each connectivity gradient separately. Positive correlations suggest that changes in functional connectivity are more pronounced in voxels that are close to the infarct region in the corresponding gradient.

For a detailed description of the analysis steps see Supplementary Figure S2.

### 2.10 The relationship between changes in functional connectivity over time and anatomical lesion location

Euclidean distances from each voxel to the infarct area in MNI152 (3 mm^3^) space using three-dimensional voxel coordinates were computed for each patient. The resulting anatomical distance values were correlated with concordance values (using Pearson’s correlation coefficient). A regression analysis was applied to remove the contribution of this factor from CCC-values. Residuals were correlated with gradient-based distance-to-lesion values (using Spearman’s rank-order correlation coefficient).

### 2.11 The relationship between changes in functional connectivity along connectivity gradients and changes in clinical scores

Individual gradients were divided into uniform parcels (bins). We varied the number of bins used for the parcellation from 5 to 3000 in order to consider the continuous nature of connectivity gradients while allowing us to classify parts of the gradients as affected by the lesion. At each bin number and for each stroke patient, bins that overlapped with lesioned-voxels were identified as “lesion-affected”, whereas the remaining bins were defined as “lesion-unaffected”. An overall delta-concordance measure, Δ*CCC*, was computed as the difference between average concordances in lesion-unaffected and lesion-affected bins, such that Δ*CCC*= µ_*unaffected*_-µ_*affected*_. A positive Δ*CCC* score reflects a higher functional connectivity alteration over time in affected bins. Of note is that lesioned voxels were removed from this computation, thereby the difference in concordance reflects the degree of preferential change in functional connectivity in affected yet structurally intact areas.

To explore the link between changes in clinical scores and the overall delta-concordance measure detected along gradients, the National Institute of Health Stroke Scale (NIHSS) was used. The NIHSS values were assessed at the day of admission (day 0) and discharge (day 5). Twenty-seven patients out of 28 completed the NIHSS assessment at both time points. Patients were divided into two groups; those who changed in clinical score from day 0 to day 5 (“clinical change”, n = 16), and those who did not change (“no clinical change”, n = 11).

Permutation test (with 10,000 iterations) was used to examine the significance of the difference in mean Δ*CCC* values for the two groups of patients (“clinical change” versus “no clinical change”). The test was repeated for each variation of bin numbers as well as for each of the three connectivity gradients. Positive values reflect that a preferential change in concordance over affected bins is more pronounced in patients who changed their clinical score from day 0 to day 5. To control for the multiple comparison problem resulting from varying the number of bins (N= 2996 tests), the False Discovery Rate (FDR) correction (Benjamini and Hochberg, 1995) was applied with a threshold of 0.1.

## 3.1 Results

### 3.2 Mapping stroke lesions onto connectivity gradients

To map heterogeneous lesions across our sample of patients, individualized lesion masks were delineated and projected onto a standard MNI brain (Figure 3A), as well as onto the first three connectivity gradients (Figure 3B). Lesions were heterogeneous in both location and size (mean volume=4.11 cm^3^, SD=2.80 cm^3^), and distributed in subcortical (n=13), cortical (n=14), and brainstem (n=1) regions. For further details on individual lesion location and affected vascular territories, see Supplementary Table 1.

**Figure 3.**
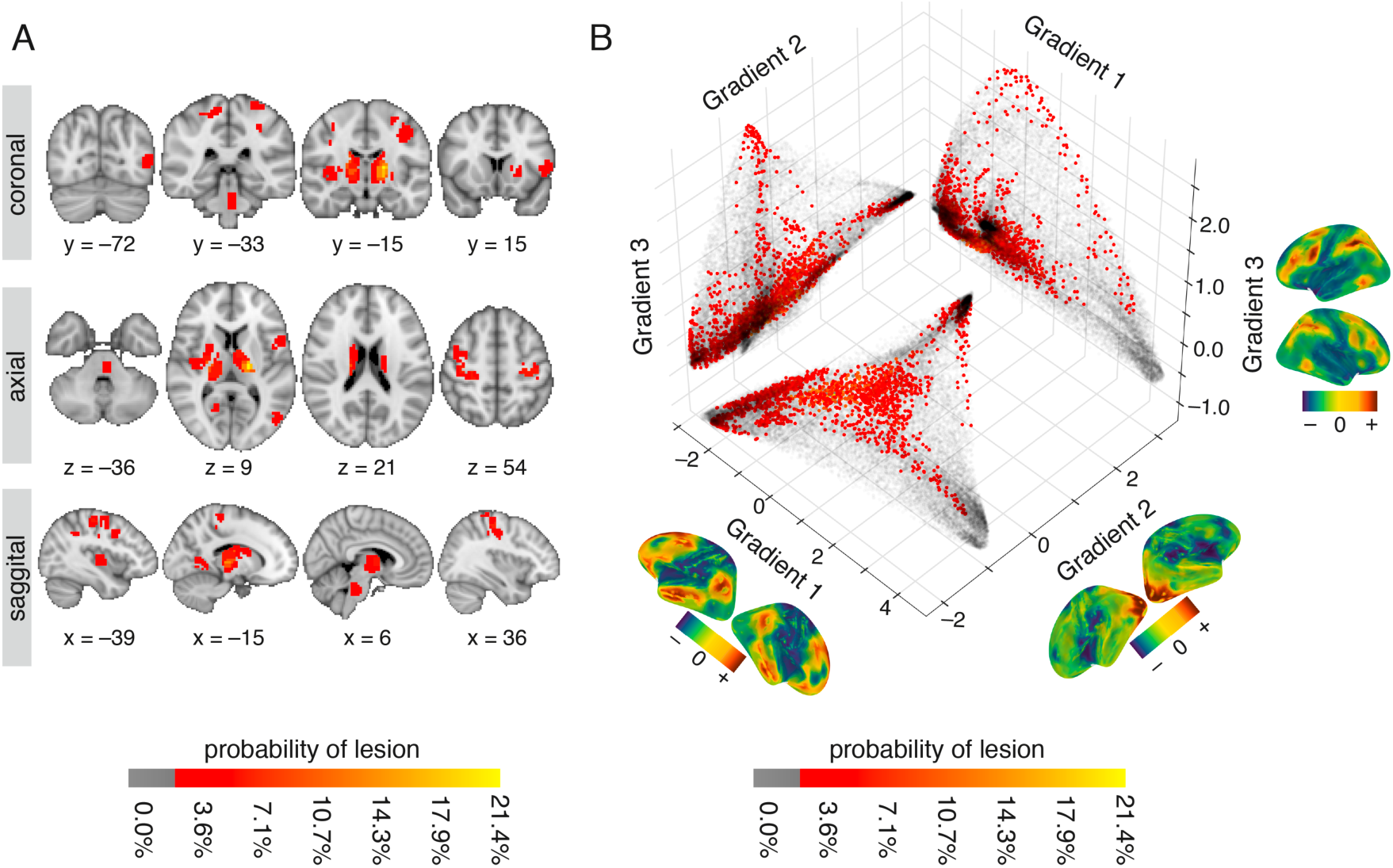
Lesion location across patients shown in anatomical space and along connectivity gradients. (A) Anatomical lesion distribution in individual stroke patients (n=28) projected onto an MNI brain. The red-to-yellow color bar indicates the percentage of patients with lesions in that voxel. (B) Location of lesions projected onto the first three connectivity gradients. The three connectivity gradients represent a low-dimensional description of the whole-brain connectivity matrix obtained using healthy controls’ data (n=28). Corresponding spatial maps of each connectivity gradient are projected on brain surface mesh near respective axes. Colors represent positive (sienna) and negative (dark blue) embedding values, in accordance with values along the axes. Along each gradient, voxels that share similar connectivity patterns are situated close to one another and have similar embedding values. Grey scatter plots depict a two-dimensional connectivity space created as a combination of any two given gradients. Lesion location along each gradient is projected onto the two-dimensional space as an alternative approach to anatomical lesion mapping. The red-to-yellow color bars indicates the percentage of patients with lesions in that voxel. Lesioned voxels are mostly clustered around the edges of the connectivity gradients such that they affect sensorimotor areas and ventral and dorsal areas associated with attention.

Projecting lesion locations onto the connectivity gradients enabled us to assess which portions of connectivity space were affected by the stroke. The template connectivity space was based on a decomposition of voxelwise functional connectivity data from healthy controls. Voxels that share functional connectivity patterns are situated closer to one another along a given connectivity gradient. For example, voxels that are part of the default-mode network are clustered at the high end of Gradient 1, and those that are part of primary sensory areas at the low end (Margulies et al., 2016). Here, we used the first three gradients that account for a total variance of 50.84% in the healthy control connectivity data (see Supplementary Figure S3).

Figure 3B demonstrates the distribution of lesioned voxels within the three-dimensional connectivity space. We found that although the anatomical location of lesions was heterogeneous (Figure 3A), within the connectivity space lesions were predominantly clustered at the extremes of each gradient, especially those of Gradients 1 and 3 (Figure 3B).

### 3.3 The impact of lesion location along specific connectivity gradients on reorganization

To determine if the location of lesions along specific gradients is associated with changes in functional connectivity after stroke, we computed for each voxel: 1) spatial concordance, which reflected the degree of change in the functional connectivity pattern over time. Spatial concordance values range between -1 and 1 such that lower values reflect a larger change in functional connectivity pattern over time; and, 2) distance-to-lesion along each connectivity gradient. Distance values represent the similarity of functional connectivity patterns for any given voxel with the lesioned area. Low distance values reflect voxels that share similar functional connectivity pattern with the lesion site. Importantly, the lesioned voxels were excluded from both these analyses such that only the indirect effects of the lesion (i.e., diaschisis) were assessed. Spatial concordance and distance-to-lesion were correlated for individual patients, and individual connectivity gradients.

We found a significant relationship between the degree of functional connectivity alterations over time and proximity of non-lesioned voxels to lesion locations along Gradient 1 and Gradient 3. No significant relationship was found for Gradient 2 (Figure 4A, Table 1).

**Figure 4.**
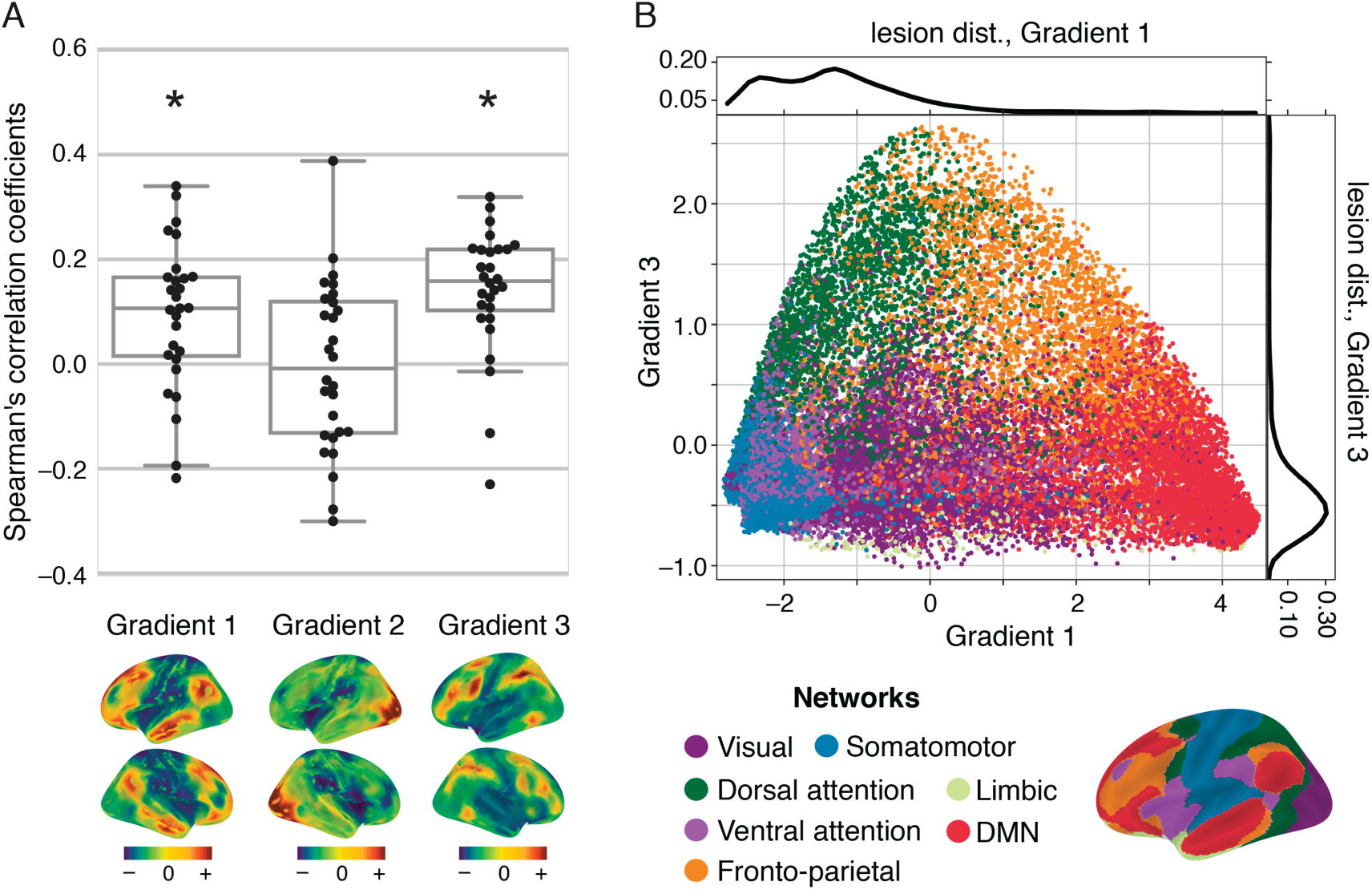
The relationship between lesion location along connectivity gradients and the degree of changes in functional connectivity in non-lesioned voxels over time. (A) Correlation values between distance-to-lesion and spatial concordance (y-axis) are shown for individual patients and the three connectivity gradients (x-axis). The spatial map of each connectivity gradient is shown below the respective location on the x-axis. Correlations were significantly positive for Gradient 1 (P=0.0027, W=71.0, one-tailed Wilcoxon signed-rank test) and Gradient 3 (P=0.0001, W=35.0), but not for Gradient 2 (P=0.76, W=189.0). The closer a voxel is to the lesioned site mapped on connectivity gradients 1 and 3, the more pronounced its functional connectivity changes over time. (B) Continuous connectivity gradients and corresponding seven canonical resting-state networks (Thomas Yeo et al., 2011). Voxels are situated based on their embedding values along Gradient 1 (x-axis) and 3 (y-axis) and colored according to their network assignment. Gradient 1 captures the dissociation between the default-mode network (DMN) and the sensorimotor networks on its two edges, while Gradient 3 captures the dissociation between dorsal attention/fronto-parietal networks and sensorimotor/DMN networks on its two edges. Lesion distributions along connectivity gradients are overlaid on the individual gradient axes. Lesions overlap most frequently with the lowest ends of Gradients 1 and 3.

**Table 1:** summary of statistical results. W; Wilcoxon signed-rank test.

|  | Gradient 1 | Gradient 2 | Gradient 3 |
| --- | --- | --- | --- |
| **r-values** | [-0.22, 0.34] | [-0.30, 0.39] | [-0.23, 0.32] |
| **median** | 0.11 | -0.01 | 0.16 |
| **W** | 71.00 | 189.00 | 35.00 |
| **p-values** | 0.0027* | 0.76 | 0.0001* |

Figure 4B demonstrates the correspondence between the connectivity space described by Gradients 1 and 3, and a canonical set of seven resting-state networks (Yeo et al, 2011). Gradient 1 captures the dissociation between the default-mode network (DMN) and the sensorimotor/visual networks, while Gradient 3 captures the dissociation between dorsal attention/fronto-parietal networks and sensorimotor/visual/DMN networks. For a descriptive analysis of the relationship between connectivity gradients and cognitive functions see Supplementary Material M3 and Supplementary Figure S4.

Given the expected partial correlation between distance from the lesion in connectivity space and anatomical distance, we further assessed whether anatomical location contributed to the relationship with connectivity space. We found a significant relationship between distance from the lesion in anatomical space and changes in functional connectivity over time (P = 0.0042, one-tailed Wilcoxon signed-rank test). However, using anatomical distance as a regressor of no interest did not alter the significance of our main result (see Supplementary Figure S5). Functional connectivity therefore preferentially changes after stroke in voxels that are proximal to the lesion location along Gradients 1 and 3. This relationship cannot be solely explained by the anatomical distance from the lesion.

### 3.4 Clinical relevance of functional connectivity alterations detected along connectivity gradients

Previous studies have linked alterations in functional connectivity with clinical trajectory (He et al., 2007; Ovadia-Caro et al., 2013; Park et al., 2011; Ramsey et al., 2016; van Meer et al., 2010), thereby supporting the functional significance of connectivity changes after stroke. We thus explored the relationship between functional connectivity changes and patients’ clinical trajectory for each connectivity gradient.

We tested for a group difference in spatial concordance in affected yet structurally intact areas between patients who demonstrated a change in clinical scores from day 0 to day 5 and those who did not. A positive difference in the mean of the two groups reflects an association between preferential changes in functional connectivity in affected areas and a change in clinical scores over the first week after stroke. To maintain the continuous nature of connectivity gradients, we varied the number of bins used to divide the gradients into parcels of equal size (bin numbers ranged from 5 to 3000). We found no significant difference between patients who changed in clinical scores and those who did not for any of the connectivity gradients, across different bin numbers. The averaged difference in mean for the two groups was 0.0014 (range: -0.004 to 0.015) for Gradient 1, 0.0095 (range: 0.003 to 0.015) for Gradient 2, and 0.011 (range: 0.0012 – 0.019) for Gradient 3. The range of corresponding p-values was 0.15 to 0.61 for Gradient 1, 0.12 to 0.4 for Gradient 2, and 0.03 to 0.46 for Gradient 3 (see Supplementary Figure S6).

## 4.1 Discussion

We found that stroke induces a gradual change in functional connectivity along specific connectivity gradients. Beginning with data acquired on the day of symptom onset, we showed that the degree of reorganization over the first week is influenced by the lesion location along connectivity Gradients 1 and 3. Voxels that are close to the lesion within this connectivity space demonstrate a preferential change in functional connectivity over time, regardless of their anatomical distance from the lesion.

We have implemented a decomposition approach that overcomes the necessity to parcellate the brain into discrete networks, retains information from single voxels and provides a data-driven template for studying reorganization at the connectome-level. We therefore show that strokes result in widespread connectivity changes that progress gradually along the connectome.

Our results are in line with previous studies that have used a priori defined networks. Functional connectivity alterations after stroke have been reported for sensorimotor, language and attention networks (Baldassarre et al., 2014; Carter et al., 2010; He et al., 2007; Ovadia-Caro et al., 2013; Siegel et al., 2016; Wang et al., 2010; Warren et al., 2009). These previous studies support the notion that localized lesions induce widespread effects in structurally intact areas connected to the lesion, creating a *diaschisis* effect (Andrews, 1991; Carrera and Tononi, 2014). Stroke is therefore not a strictly localized pathology (Corbetta, 2010; Ovadia-Caro et al., 2014; Ward, 2005). Remote, structurally intact areas undergo functional changes as part of the reorganization process.

Here, we extend these findings to the continuous representation of the connectome. We demonstrate that reorganization, as reflected in functional connectivity alterations, changes as a function of the distance along specific connectivity gradients. However, it is not exclusively restricted to the affected network. Thus, while most pronounced changes take place in connected areas, the effects of stroke gradually spread along the connectome.

We found that connectivity Gradients 1 and 3 better predicted the impact of a lesion on functional connectivity than Gradient 2. The three connectivity gradients capture distinct connectivity axes, with different functional domains on their extremes. One crucial difference between these gradients is that Gradient 2, in contrast to the others, represents a spectrum of relatively local patterns of connectivity (Felleman and Van Essen, n.d.; Markov et al., 2014), spanning sensory and motor systems. Regions emphasized in Gradient 2 are less likely to demonstrate changes following localized lesions, as there is little redundancy owing to long-distance connectivity. However, it remains to be investigated if changes in functional connectivity can be captured along Gradient 2 using a more homogenous lesion sample impacting only the far extremes of this gradient.

Our study demonstrates the importance of the lesion location within connectivity space for understanding the reorganization of functional connectivity. However, distance from the lesion in connectivity space is partially related to the anatomical distance, as areas close to one another often have similar connectivity patterns. In addition, local physiological changes in areas directly surrounding the lesion (Dirnagl et al., 1999) can also contribute to changes in functional connectivity (Khalil et al., 2017; Siegel et al., 2016). We therefore calculated in a control analysis the Euclidian distances from each voxel to the infarct area using a three-dimensional anatomical space. We found a significant relationship between distance based on anatomy and changes in functional connectivity as measured by spatial concordance. However, when regressing out the contribution of this factor from our main analysis, the results did not change (see Supplementary Figure S5). Consequently, changes in functional connectivity detected along connectivity gradients could not be solely explained by lesion topography or physiological processes occurring in the vicinity of the lesion site. In addition, this analysis emphasizes the significant contribution of functional connectivity changes in distant areas to the global process of reorganization.

The link between changes in functional connectivity after stroke, clinical deficits and clinical recovery has been previously shown (He et al., 2007; Ovadia-Caro et al., 2013; Park et al., 2011; Ramsey et al., 2016; van Meer et al., 2010). Here, we applied an exploratory analysis of the relationship between lesion location along connectivity gradients, changes in functional connectivity, and changes in clinical scores (NIHSS) over the first week. We divided the patients into two groups according to whether or not a clinical change took place over the first week.

Given previous findings, we expected a significant difference between the groups in the degree of change in functional connectivity patterns, however, we found no such difference for any of the connectivity gradients. Of interest nevertheless is that for Gradient 2 and Gradient 3, group differences were not randomly distributed and were positive in values (see Supplementary Figure S6).

The lack of a relationship between changes in functional connectivity and changes in clinical scores could be explained by the usage of NIHSS. NIHSS is the most commonly used assessment scale in routine acute stroke management. However, this score is fairly coarse and is not designed to accurately detect individual neurological deficits. It is instead intended to provide a standardized and reproducible overall assessment of how stroke affects a patient’s neurological status (Lyden, 2017). The relationship between functional connectivity changes along specific connectivity gradients and stroke symptomology assessed using a more detailed clinical assessment (which would better fit the voxelwise information retained in the gradients, particularly for parcellations that contain a small number of voxels) remains to be investigated in a larger sample of patients.

The conceptual shift from mapping brain regions to networks has provided a substantial improvement in how we understand the organization of functional systems. Here we aimed to translate the recent descriptions of a low-dimensional connectivity space to the clinical question of stroke-induced damage. While future studies will be necessary to better understand the utility of this framework for stroke prognosis, the current findings provide support for conceptualizing brain connectivity within a continuous connectivity-defined space. Brain networks describe interconnected regions, but similar to the problem of lesion delineation, they also require the delineation of discrete boundaries. Connectivity space offers an advance by representing the continuous nature of brain networks, but also by capturing their relative similarity. Further work is necessary to develop a mode of describing this space in a cognitive and clinical neuroscience context. Nevertheless, the current findings demonstrate its utility for capturing the impact of localized damage to the space.

## 5.1 Conclusions

Studying changes in functional connectivity after stroke in a longitudinal manner provides insight into the process of reorganization during the recovery of function. Connectivity gradients represent a methodological advancement in how we depict functionally meaningful information in the connectome. Using this fine-grained template that considers all connections has the potential of informing more targeted stroke therapies that have yet to translate to clinical usage, mostly due to oversimplified models of brain reorganization (Di Pino et al., 2014).

## Supporting information

## Acknowledgements

The authors would like to thank the patients for participating in the study, and to Dr. Julia Huntenburg, Dr. Luke Tudge and Sabine Oligschläger for their advice in various stages of this project.

