## Supplementary material for "The impact of ischemic stroke on connectivity gradients"

### M1. Quality assessment

Subjects with corrupted raw MRI data (n=4 patients) were excluded from the analysis. The image quality of resting-state MRI scans was assessed by visual inspection of the temporal signal-to-noise ratio (tSNR) distribution and mean time series across slices. Subjects displaying EPI distortion effect (n=1 control) were excluded from the analysis. Functional images exhibiting high degree of head framewise displacement (FD > 1 mm) in more than 15 volumes were discarded (n=1 patient) (Power et al., 2012). Based on the mean FD-value distribution across all scans, images with a mean head displacement exceeding 3.5 mm were excluded (n=1 control, n=5 patients). The impact of the motion control was demonstrated with whole-brain signal measures before and after motion correction. This included measuring the mean square root of differentiated time series of each voxel and the global BOLD signal (Power et al., 2014). Image co-registration quality was visually inspected with the FreeSurfer's automated white matter contours and mean functional image overlays. In cases of poor alignment between white matter contours and mean EPI images, subjects were excluded from further analysis (n=1 control). The source code for the automated quality assessment pipeline is available at https://github.com/sheyma/mriqc/tree/master, and an example of a quality assessment report at https://github.com/sheyma/mriqc/blob/master/mriqc/examples/sd02_day1.pdf.

| ID | vascular stroke territory | age | sex | lesion volume ($cm^3$) | scan days post-stroke | NIHSS admission | NIHSS discharge | $r_S$ (G1) | $r_S$ (G2) | $r_S$ (G3) |
|---|---|---|---|---|---|---|---|---|---|---|
| sd02 | L, AchA | 47 | m | 5.18 | days 0, 1, 5 | 6 | 1 | 0.16 | -0.28 | 0.27 |
| sd05 | L, MCA | 51 | m | 7.48 | days 0, 1, 5 | 5 | 3 | 0.14 | -0.06 | 0.30 |
| sd08 | L, MCA | 64 | m | 4.08 | days 0, 1, 5 | 1 | 0 | -0.06 | 0.15 | -0.13 |
| sd10 | L, MCA | 67 | f | 3.65 | days 0, 1, 5 | 0 | 0 | -0.01 | 0.04 | 0.09 |
| sd13 | R, MCA | 76 | m | 1.86 | days 0, 1, 5 | 7 | 0 | 0.04 | -0.03 | 0.17 |
| sd14 | L, MCA | 72 | f | 1.22 | days 0, 1, 5 | 0 | 0 | 0.11 | -0.13 | 0.15 |
| sd16 | R, MCA | 54 | m | 2.03 | days 0, 1, 5 | 1 | 0 | -0.19 | 0.13 | 0.11 |
| sd17 | L, PCA | 43 | m | 2.11 | days 0, 1, 5 | 3 | 2 | 0.09 | 0.39 | 0.07 |
| sd21 | R, MCA | 72 | f | 14.09 | days 0, 1, 5 | 1 | 0 | 0.07 | 0.09 | 0.18 |
| sd25 | L, PCA | 71 | f | 6.80 | days 0, 1, 5 | 5 | 0 | 0.10 | 0.12 | 0.18 |
| sd26 | L, PCA | 79 | m | 3.19 | days 0, 1, 5 | 2 | - | 0.14 | -0.04 | 0.16 |
| sd28 | R, PCA | 73 | f | 3.38 | days 0, 1, 5 | 5 | 2 | 0.32 | -0.17 | 0.22 |
| sd31 | R, PCA | 81 | f | 3.21 | days 0, 1, 5 | 4 | 2 | 0.17 | -0.10 | 0.21 |
| sd32 | L, MCA | 47 | m | 2.51 | days 0, 1, 5 | 11 | 0 | -0.06 | -0.30 | 0.01 |
| sd33 | L, PCA | 68 | m | 4.37 | days 0, 1, 5 | 1 | 1 | 0.34 | 0.16 | 0.13 |
| sd35 | L, MCA | 75 | f | 7.26 | days 0, 1, 5 | 3 | 2 | 0.25 | 0.03 | 0.22 |
| sd36 | L, MCA | 50 | m | 2.27 | days 0, 1, 5 | 1 | 1 | 0.27 | -0.14 | 0.32 |
| sd38 | R, PCA | 53 | m | 1.46 | days 0, 1, 6 | 1 | 0 | 0.02 | 0.12 | 0.22 |
| sd39 | L, PCA | 59 | m | 4.75 | days 0, 1, 5 | 1 | 1 | 0.02 | 0.09 | 0.15 |
| sd40 | R, PCA | 90 | f | 7.61 | days 0, 1, 5 | 6 | 2 | 0.01 | 0.20 | -0.01 |
| sd42 | R, MCA | 72 | m | 3.89 | days 0, 1, 5 | 0 | 0 | -0.11 | -0.13 | 0.09 |
| sd43 | R, AchA | 75 | f | 2.51 | days 0, 1, 5 | 4 | 4 | 0.13 | 0.01 | 0.13 |
| sd44 | R, MCA | 78 | m | 3.43 | days 0, 1, 5 | 1 | 2 | 0.17 | -0.14 | 0.23 |
| sd45 | L, MCA | 37 | f | 1.78 | days 0, 1, 4 | 0 | 0 | -0.22 | -0.22 | -0.23 |
| sd46 | L, MCA | 63 | f | 5.43 | days 0, 1, 4 | 1 | 2 | 0.18 | -0.05 | 0.25 |
| sd48 | L, PCA | 60 | m | 0.73 | days 0, 1, 4 | 1 | 1 | 0.16 | -0.17 | 0.14 |
| sd49 | R, PCA | 63 | m | 6.75 | days 0, 1, 5 | 1 | 1 | 0.11 | 0.10 | 0.11 |
| sd52 | R, MCA | 81 | m | 2.13 | days 0, 1, 5 | 0 | 0 | 0.25 | 0.17 | 0.22 |
| | **Mean** | 65.04 | | 4.11 | | 2.57 | 1.00 | 0.09 | -0.01 | 0.14 |
| | **std** | 13.27 | | 2.80 | | 2.70 | 1.09 | 0.14 | 0.16 | 0.12 |

**Supplementary Table 1: Patient characteristics and statistical results.**
Abbreviations: AChA indicates Anterior Choroidal Artery; PCA, Posterior Cerebral Artery; MCA, Middle Cerebral Artery; L, left; R, right; B, bilateral; f, female; m, male. NIHSS (National Institute of Health Stroke Scale) scores were obtained at patient admission and discharge days, $r_S$: Spearman's rank-order correlation coefficient between concordance and distance-to-lesion maps along Gradient 1 (G1), Gradient 2 (G2) and Gradient 3 (G3).

#### M2. Controlling for variation in the mask size

Since the grey matter masks used for the patients were computed at the individual level, they varied slightly among patients (mask size range: 32,659 - 33,212 voxels). In order to make sure this variation had no influence on our main result, we repeated the analysis using a common mask (containing 30,314 voxels prior to lesion removal).

An averaged whole-brain resting-state mask was obtained across all stroke patients and scan time points. This mask was multiplied with the grey matter template of healthy controls, resulting in a new grey matter template of 30,134 voxels. The dilated lesions were excluded from the group-level grey matter template for individual patients. Using this more restricted grey matter template had no influence on the main result (Supplementary Figure S1).

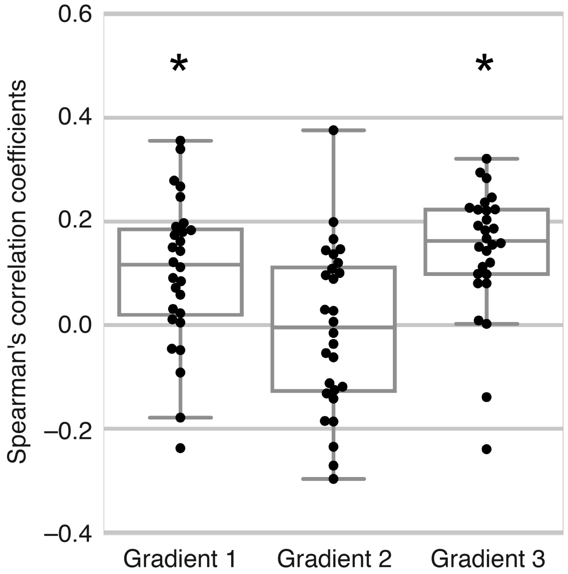


**Supplementary Figure S1: Control analysis to overcome voxel number variation across patient's specific grey matter masks.** Our main result is replicated using a common grey matter template that was obtained across all stroke patients and resting-state sessions. Correlations were significantly positive for Gradient 1 (P=0.0015, W=63.0, one-tailed Wilcoxon signed-rank test) and Gradient 3 (P=0.0001, W=33.0), but not for Gradient 2 (P=0.76, W=189.0). Median correlations are $\tilde{r}$=0.12 for Gradient 1, $\tilde{r}$=0.00 for Gradient 2, and $\tilde{r}$=0.16 for Gradient 3.

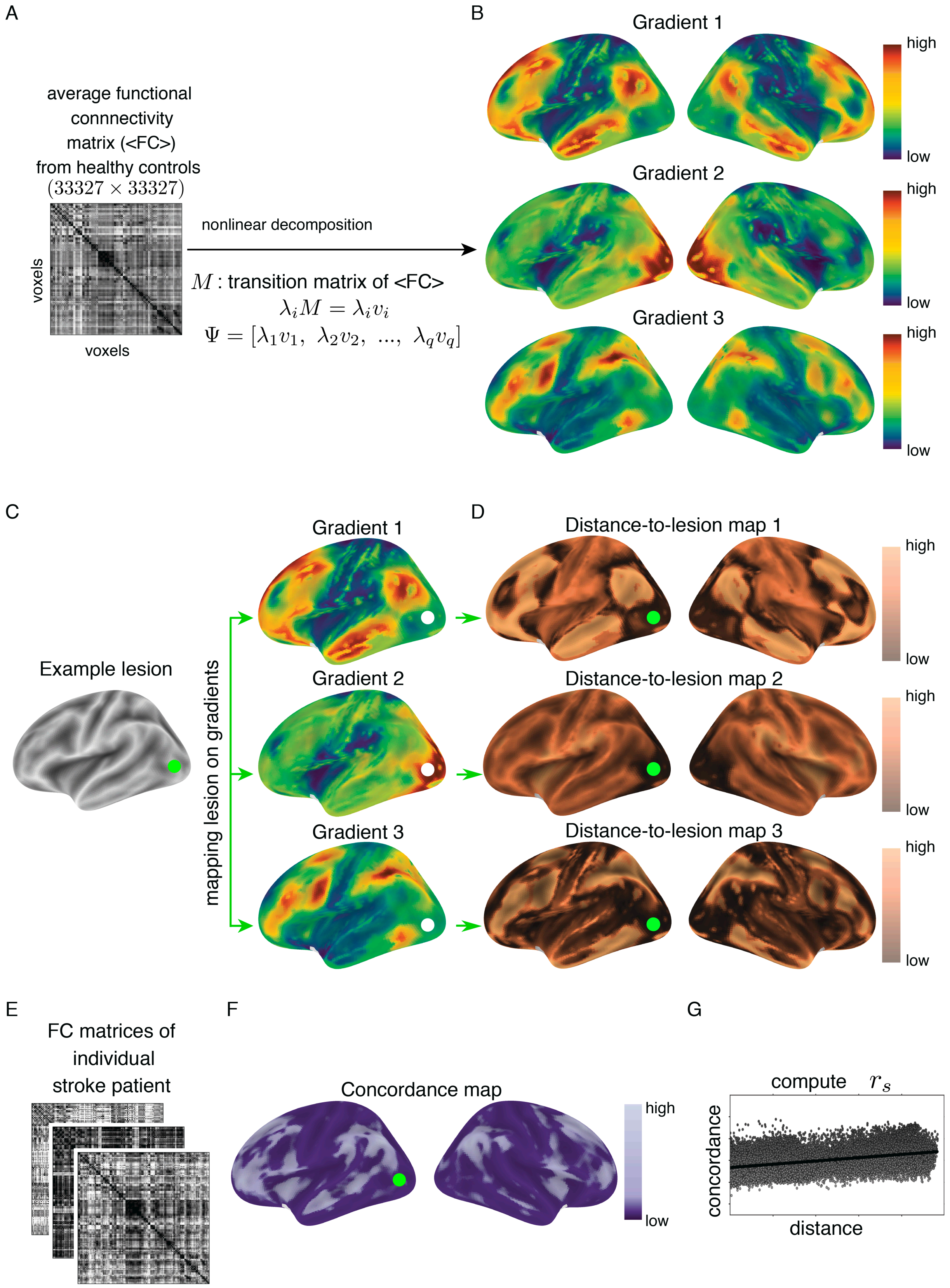


**Supplementary Figure S2: A detailed description of the analysis pipeline.** (A) A schematic illustration of averaged voxelwise functional connectivity matrix (<FC>) based on resting-state fMRI data of healthy subjects (n=28). (B) Applying a nonlinear decomposition analysis on healthy controls data reveals first three connectivity gradients. (C) Anatomical lesion locations were delineated for individual patients and located along gradients. (D) The distance from each voxel and the lesioned site was computed to create a voxelwise distance-to-lesion map for each gradient. Distance-to-lesion is the subtraction of embedding values between each voxel and the mean lesioned voxels. Voxels with lower values on the distance map (dark copper) shared similar functional connectivity patterns with the lesioned site as characterized in healthy controls. (E) For each patient, voxelwise functional connectivity (FC) matrices were computed for the three resting-state fMRI scans post-stroke. (F) Connectivity concordance correlation coefficient was used to quantify changes in functional connectivity patterns over time. Lower concordance values (dark purple) reflect a larger change in functional connectivity patterns over time. (G) For each gradient and each individual patient, the relationship between the distance-to-lesion and concordance values was tested using Spearman's rank-order correlation coefficient ($r_s$) at the voxelwise level. The lesioned voxels were excluded from this analysis to capture indirect, rather than local effects of the lesion.

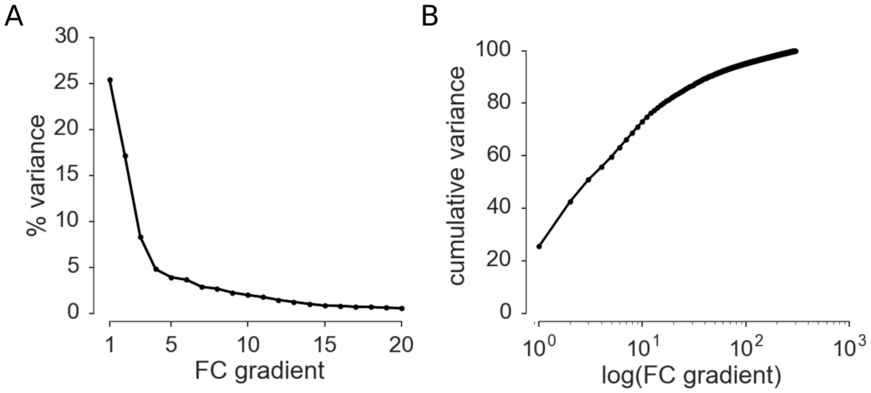


**Supplementary Figure S3: Variance explained by connectivity gradients based on nonlinear decomposition of functional connectivity data from healthy controls (n=28).** The variance percentage (y-axis, A) explained by the first 20 gradients (x-axis, A) and the cumulative variance (y-axis, B) carried by the first 300 gradients (x-axis, B) are depicted. The variance drops exponentially with increasing number of gradients. The first three gradients used for the current analysis account for 50.84 % of the total variance.

#### M3. The distribution of cognitive functions along connectivity gradients

Previous studies have demonstrated the cognitive dissociation captured along Gradient 1 (Margulies et al., 2016). However, this descriptive relationship has not been shown for Gradient 3. We therefore conducted a meta-analysis using the NeuroSynth database (Yarkoni et al., 2011) to assess the topic terms associated with location along Gradient 3. Analysis followed a previously published procedure (Margulies et al., 2016), and included 24 topic terms (Figure S4). The volumetric map of Gradient 3 was divided into bins of five percentile size, creating 20 maps/regions-of-interest ranging from 0-5% to 95-100%. Each of the 20 maps was then binarized and fed as an input to the meta-analysis. The output of the analysis was a z-statistic associated with the feature term for each given map. The terms were ordered based on the weighted mean for visualization.

This descriptive analysis supports the functional dissociation of transmodal (i.e., default-mode network, DMN) and unimodal areas from multi-modal regions as captured along Gradient 3.

Topic terms such as ‘social cognition’ associated with DMN activation are situated at the far end of the gradient, followed by unimodal areas associated with topic terms such as ‘auditory’ and ‘motor’. At the other extreme of the gradient, topic terms such as ‘visuospatial’ and ‘working memory’ are associated with areas of the attention and memory domains.

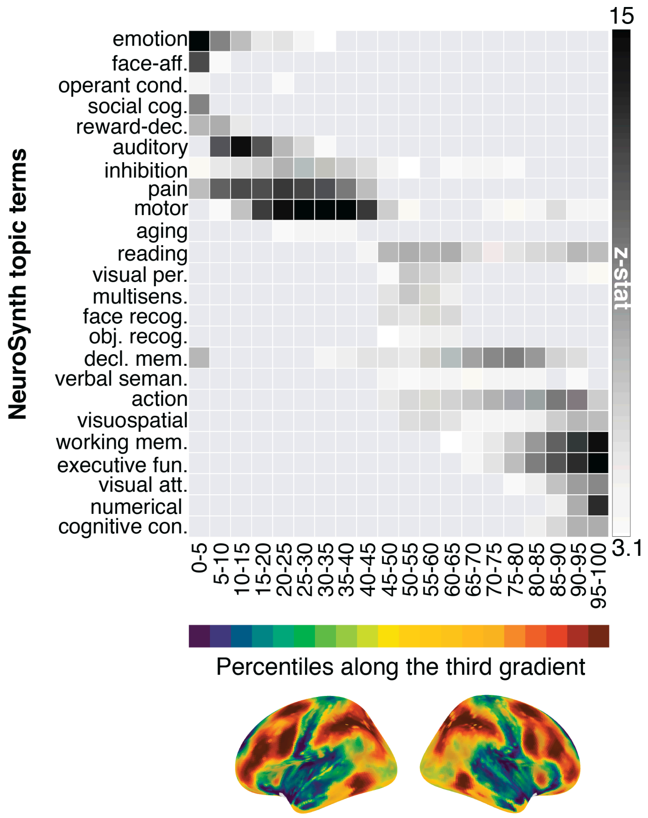


**Supplementary Figure S4: NeuroSynth meta-analysis of regions of interest along Gradient 3 using 24 topic terms.** The gradient is divided into 20 bins of equal size each representing five percentiles (blue-to-red colorbar and corresponding spatial map of the gradient). The z-statistic (grey colorbar) represents the association with a topic term for a given map. Topic terms are ordered by the weighted mean of their location along the gradient. High-order abstract terms associated with default-mode network at the top are followed by sensory processing terms, and then at the far end memory and attention related processes. *Abbreviations: face-aff.: face-affective processing; operant cond.: operant conditioning; social cond.: social conditioning: reward-dec.: reward-based decision making; auditory: auditory processing; visual per.: visual perception; multisens.: multisensory processing; face recog.: face recognition; obj. recog.: object recognition; decl. mem.: declarative memory; verbal seman.: verbal semantics; working mem.: working memory, executive fun.: executive function, visual att.: visual attention; numerical: numerical cognition; cognitive con.: cognitive control.*

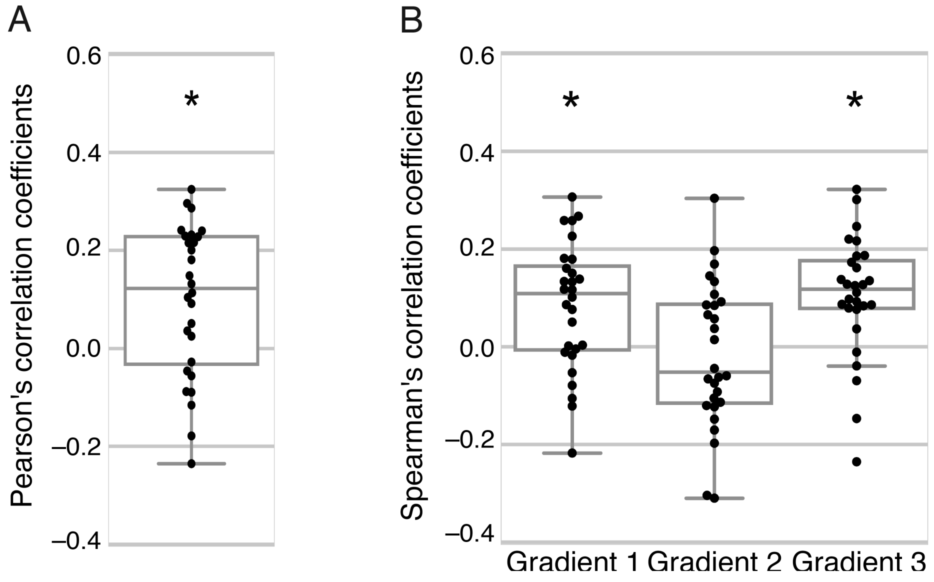


**Supplementary Figure S5: The influence of anatomical lesion location on functional connectivity changes.** (A) Box plot indicates the distribution of Pearson's correlation coefficients for individual patients (y-axis) representing the relationship between distance from the lesion in anatomical space and changes in functional connectivity over time. A positive correlation indicates that voxels situated nearby the stroke lesion have a higher alteration in their functional connectivity patterns. One-tailed Wilcoxon signed-rank test shows a significant positive correlation at the group level ($\tilde{r}=0.12$, $P=0.0042$, $W=77.0$). Functional connectivity alters at a higher degree for voxels that are anatomically closer to the lesion. (B) Box plots represent the distribution of Spearman's rank-order correlation coefficients from individual patients (y-axis) for three main gradients, respectively (x-axis) following the regression of anatomical distance as a covariate of no interest. A positive correlation reflects a significant relationship between changes in functional connectivity and location on respective connectivity gradients independent of the anatomical lesion location. The relationship between location along connectivity gradients and functional connectivity changes remains unchanged for all three gradients (one-tailed Wilcoxon signed-rank test, Gradient 1: $\tilde{r}=0.11$, $P=0.0046$, $W=78.0$, Gradient 2: $\tilde{r}=-0.05$, $P=0.50$, $W=173.0$, Gradient 3: $\tilde{r}=0.12$, $P=0.0006$, $W=51.0$).

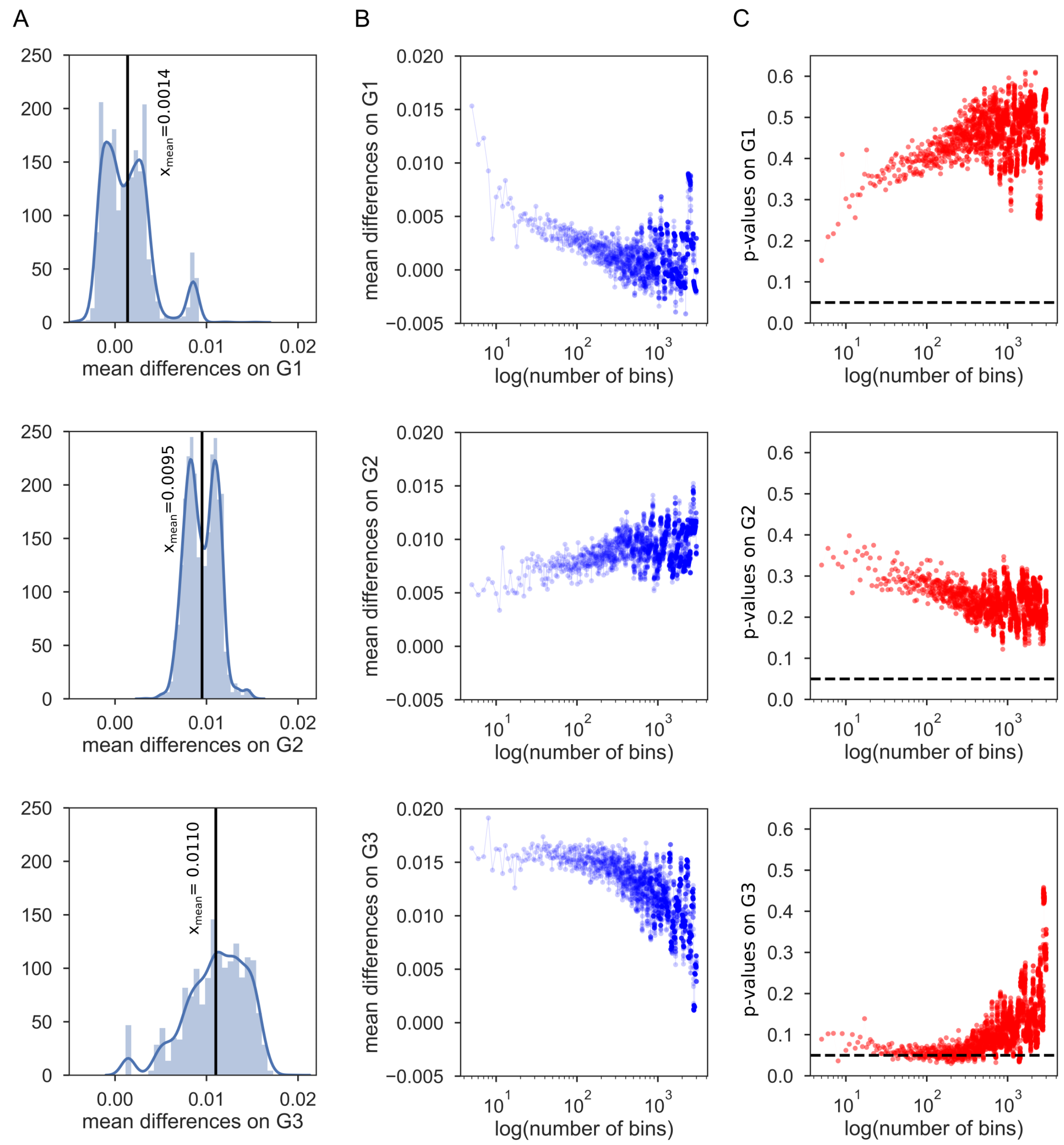


**Supplementary Figure S6: Relationship between the clinical trajectory and overall delta-concordance measure detected along individual gradients.** (A) Distribution of the difference in mean delta-concordance ($\Delta CCC$) between patient groups ("clinically-changed" versus "no clinical change") across varying bin numbers (from 5 to 3000) are plotted for individual gradients (Gradient 1, top; Gradient 2, middle; Gradient 3, bottom). $\Delta CCC$ scores depict the spatial concordance differences in lesion-unaffected and lesion-affected bins of a gradient. The difference in mean $\Delta CCC$ between patient groups was computed for each of the bin numbers used to parcellate the connectivity gradient into parcels of equal size. A positive difference suggests that the group of patients who changed their clinical score over the first week demonstrated a higher degree of change in functional connectivity in affected portions of the gradient. We found no significant difference between the groups across varying bin numbers following False Discovery Rate (FDR) correction (threshold = 0.1, number of tests = 2996). Values are centered around zero for Gradient 1, however, they are positively distributed for

Gradient 2 and Gradient 3. (B) The mean difference in $\Delta CCC$ between patient groups (y-axis) depicted for each bin number (x-axis). (C) The corresponding p-values (y-axis) for each bin number (x-axis) obtained following group comparison using a permutation test. The dashed black line represents $p = 0.05$.
